# Machine learning approaches to predicting whether muscles can be elicited via TMS

**DOI:** 10.1101/2023.12.14.571623

**Authors:** Fang Jin, Sjoerd M. Bruijn, Andreas Daffertshofer

## Abstract

**Background:** Transcranial magnetic stimulation (TMS) is a valuable technique for assessing the function of the motor cortex and cortico-muscular pathways. TMS activates the motoneurons in the cortex, and this activation is transmitted through the cortico-muscular pathway, after which it can be measured as a motor evoked potential (MEP) in the muscles. The position and orientation of the TMS coil and the intensity used to deliver a TMS pulse are considered central TMS setup parameters influencing the presence/absence of MEPs.

**New Method:** We sought to predict the presence of MEPs from TMS setup parameters using machine learning. We trained different machine learners using either within-subject or between-subject designs.

**Results:** We obtained prediction accuracies of on average 77% and 65% with maxima up to up to 90% and 72% within and between subjects, respectively. Across the board, a bagging ensemble appeared to be the most suitable approach to predict the presence of MEPs, although a comparably simple logistic regression model also performed well.

**Conclusions:** While the prediction between subjects clearly leaves room for improvement, the within-subject performance encourages to supplement TMS by machine learning to improve its diagnostic capacity with respect to motor impairment.

## 1. Introduction

Transcranial magnetic stimulation (TMS), a non-invasive brain stimulation technique, is a popular and well-established tool, both for experimental assessments and in the clinic (Kobayashi and Pascual-Leone, 2003). The first TMS-induced muscle activation (magnetic evoked potential, MEP) was captured in 1985 (Barker et al., 1985). The activation originated from depolarisation of the upper motoneurons due to a TMS pulse, which passed via the descending neural pathway to the muscle, where it can be recorded by conventional electromyography (EMG). The assessment of motor neurophysiological outcomes like the active area of TMS motor mapping is guided by the stimulation of sites on the (motor) cortex from which an MEP can be elicited, coined active points (Krieg et al., 2017). Quite recently, however, we showed that incorporating so-called non-active points, i.e., sites on the (motor) cortex from which no MEP can be elicited, can improve estimates of these motor neurophysiological outcomes.

The position of the TMS coil, or more precisely the position of the focal point of the induced magnetic field, is a prime parameter determining whether a certain muscle is activated. For instance, stimulation in the hand-knob region of the motor cortex can trigger finger muscles (Dechent and Frahm, 2003). Apart from position, however, coil orientation and stimulation intensity also determine whether an MEP will be elicited (Laakso et al., 2013; Opitz et al., 2013; Weise et al., 2020). Vink et al. (2018) showed that MEP amplitudes altered with changing coil orientation even when the stimulation occurred in the same position. Here we asked to what extent one can predict whether a TMS stimulation with a certain intensity and at a certain position and orientation does (or does not) elicit an MEP in a muscle. If possible, this may greatly facilitate performing TMS-based assessments of the motor cortex.

One way to achieve this would be to define a very detailed biophysical model of the neural tissue in the cortex and to estimate the induced electric field for every possible location, orientation, and intensity of the TMS stimulation. Not only will that be a laborious endeavour, but this approach will also come with unavoidable uncertainties about the neural responses to TMS. Therefore, we considered a blind alternative: Supervised machine learning uses existing data for training a model to predict the unseen output given some input. We adopt this approach to classify the presence of MEPs (= output response) from TMS setup parameters, namely position, orientation, and intensity (= input predictors).

We tested the predictive capacity of different machine learners using data for N=20 subjects in two distinct settings. (1) We adopted a within-subject design by utilising parts of the data from a single subject as a training set used to predict the remaining data from the same subject. (2) We tried to predict the presence of MEPs from an unseen subject using the data of other the subjects, that is, we considered a between-subject design. For both designs we anticipated challenges. For the within-subject case the amount of data of an individual subject may not suffice to reach maximal prediction accuracy. We return to this in the discussion section as sample size is crucial for machine learning to be successful. In the between-subject case one faces pronounced inter-individual differences in brain morphology that we sought to minimise by warping the coil position (Jin et al., 2023) and orientation from specific-subject MRI to a standard anatomical template.

If within-subject prediction is feasible, it may help streamlining motor mapping procedures for identifying the motor region in general but especially in patients prior to surgery (Krieg et al., 2017). To ensure the precision of motor mapping, more than 100 stimulation points are typically required (Van De Ruit et al., 2015). Reducing that number by any means will decrease valuable time needed otherwise in the operating theatre. The potential benefits of between-subject prediction are more abundant. For example, the active area induced by TMS is a biomarker in stroke (Sawaki et al., 2008). Using healthy controls as training set one may generate a reference area of excitability that may serve as a target when evaluating therapeutical interventions in stroke survivors.

## 2. Methods

### 2.1. Data collection

The data were collected in a previously described study (Jin et al., 2022). In brief, N=20 subjects underwent TMS motor mapping. We used a Magstim 200^2^ stimulator (Magstim Company Ltd., Whitland, Dyfed, United Kingdom) with neuro-navigation (Neural Navigator, Brain Science Tools BV, De Bilt, The Netherlands). Bipolar EMG signals (Porti, TMSi, Oldenzaal, the Netherlands) were recorded from the muscles of the first dorsal interosseous (FDI), abductor digiti minimi (ADM), flexor pollicis brevis (FPB), abductor pollicis brevis (APB), extensor digitorum communis (EDC), flexor digitorum superficialis (FDS), extensor carpi radialis (ECR), and flexor carpi radialis (FCR). Prior to the measurement, each subject underwent an anatomical T1 MRI (3 Tesla Philips Achieva System, Philips, Best, Netherlands). We mapped the motor cortex using a pseudo-random walk method (Van De Ruit et al., 2015) at intensities of 105% RMT of FDI, EDC and FCR and recorded the EMG signals simultaneously. At each of the three intensities about 120 stimulations interleaved by 5s rest were performed. All the measurement procedures were carried out twice, yielding a total of about 120·3·2 = 720 stimulation per subject.

### 2.2. Predictors and response data

We selected TMS setup parameters (intensity, position, and orientation of TMS) as the input data (predictors) and the presence/absence of MEPs as the output data (categorical response) for our machine learners. The presence of (= MEP) was defined as equal to 1 if the stimulation yielded an MEP in any of the eight muscles, whereas if none of them was elicited we considered MEPs to be absent (= no MEP). Please note that all the details about determining MEP from EMG can be found in (Jin et al., 2022). In brief, we considered MEPs proper if their amplitude exceeded 20 times the EMG-baseline’s standard deviation but stayed below 10 mV, and if the peak’s latency fell within the range of 5 to 50 ms after stimulation. All other stimulations were marked as no MEPs.

To standardise the input variables, we expressed the intensity relative to the FDI resting motor threshold (RMT) rather than in stimulator output. We also converted the Cartesian position data into spherical coordinates, i.e., (*x, y, z*) ⟶(*r, φ*, *ϑ*), where r denotes the radius and *φ* and *ϑ* the azimuth and elevation, respectively. And we expressed the rotation matrix that quantifies the coil orientation via the corresponding Euler angle (*α, β, γ*). For all angular variables we computed the corresponding sine- and cosine-coefficient to account for their 2π-periodicity. That is, we used a total of twelve input variables: intensity (in RMT units), the radius *r*, and 2·5 = 10 sine- and cosine-coefficients of the five angular variables.

### 2.3. Machine learners

We considered four types of machine learners. The first two, (1) a logistic regression model with ridge-regression for regularisation and (2) a regular support vector machine (SVM), assess whether the data sets are linearly separable, for (2) after embedding the data in a (much) higher-dimensional space (Steinwart and Christmann, 2008). The other two are non-linear approaches: (3) A support vector machine using Gaussian kernel functions when embedding the predictor data; and (4) random decision trees combined into a so-called bootstrap aggregating (bagging) ensemble model (Breiman, 1996). In a nutshell, data are subsampled yielding bootstraps with which different decision trees are trained whose outcomes are aggregated based on a majority vote. Bagging is known to reduce overfitting and appears robust against random fluctuations in training data.

For every type of machine learner, we first estimated a basic set of hyperparameters by pooling data and fitting the respective model via Bayesian optimisation and 20-fold crossvalidation; see Table A1 for the resulting sets. Subsequently we used them for the training/validation steps before testing the resulting models on unseen data.

### 2.4. Within-subject design

As said, we first pooled the data, here, of an individual subject. With that we estimated the optimal hyperparameters via 20-fold cross-validation and fixed them for the subsequent testing. To guarantee that both MEP and no MEP stimulations were properly represented, for each of the 20 folds we stratified the data accordingly. For testing the models, we used a holdout validation, i.e., we used 80% of the data for training and validation (Bayesian optimisation with 10-fold cross-validation with fixed hyperparameters, again with proper data stratification) and estimated the predictive capacity of the trained model using the held-out 20% data as test set. As illustrated in Figure 1, we repeated this for every of N=20 subjects 25 times and computed the resulting confusion chart.

**Figure 1.**
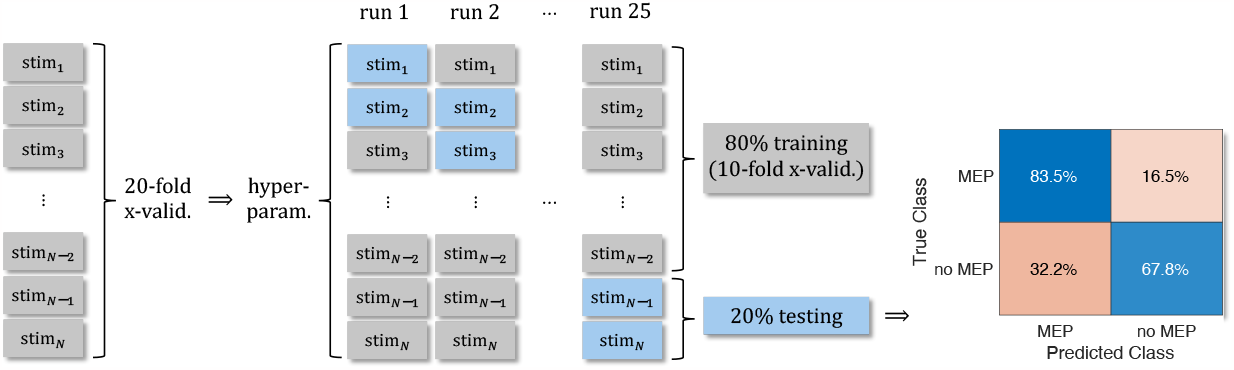
Flow chart of the within-subject assessment, from left to right: After an initial training/validation step to fix the models’ hyperparameters, we adopted a 10-fold cross-validation to train the model on 80% of the data and used the remaining 20% for testing. Twenty-five repetitions of this procedure resulted in a confusion chart illustrating how many false/true positives/negatives were predicted. In the Results section, we report the corresponding averages over N=20 subjects.

This is generally characterised via the percentage of true positives (TP), false positives (FP), false negatives (FN), and true negatives (TN). Here, the ‘true positive’ refers to predicting the presence of a MEP if a MEP had been elicited, whereas a ‘false positive’ implies predicting its absence although a MEP had been elicited. Likewise, a ‘true negative’ implies the prediction of the absence of a MEP when it had not been elicited and a ‘false negative’ refers to the prediction of a MEP in that case.

We averaged the confusion charts over 25 repetitions per subject and report the mean over subjects as main outcome. The number of correct predictions (TP+TN) divided by the number of all predictions (TP+FP+FN+TN) served as a measure of the model’s accuracy and/or predictive capacity. We chose this measure of accuracy over the overall k-fold loss, because it avoids a potential bias in case the numbers of stimulations with and without MEPs differ substantially. For example, if 80% of all stimulations elicited MEPs, then always predicting the presence of an MEP would yield a 20% k-fold loss, i.e. 80% accuracy.

### 2.4. Between-subject and within-subject design

With the between-subject models we tested for the capacity to predict the MEP/no-MEP outcome after training our machine learners using the data of all-but-one subjects. Otherwise, our approach largely corresponded to the within-subject design, i.e. we pooled the data, here of all subjects, to fix the hyperparameters of the four different types of machine learners, before training the models using the data of N-1=19 subjects and testing them on the single remaining subject; see Figure 2. We repeated this for all the twenty hold-one-out combinations and report below the average confusion chart as well as the corresponding model accuracy/predictive capacity. The results for the individual subjects are given in the Appendix.

**Figure 2.**
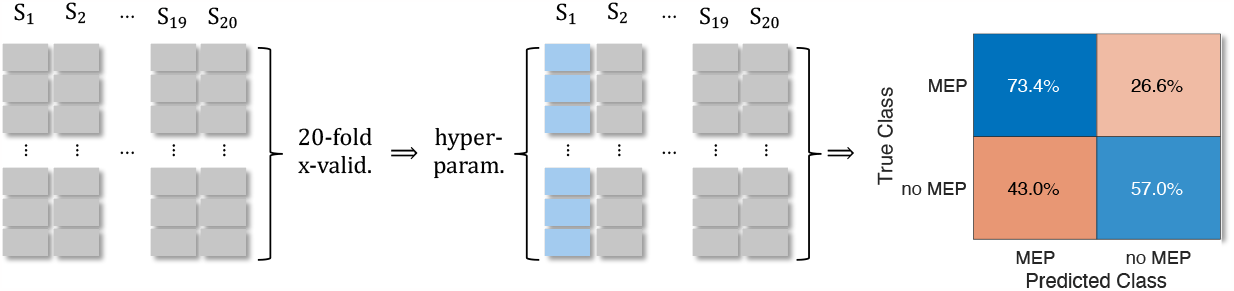
Flow chart of the between-subject assessment, from left to right: After a 20-fold cross-validation initial training/validation step to fix the models’ hyperparameters, we adopted to train the model on the data of one subject and used the remaining data of other 19 subjects for testing. This procedure was repeated for every of the 20 subjects. The resulting confusion chart illustrates how many true/false positives/negatives were predicted, e.g. how often the presences of a MEP was predicted when it had been predicted (= ‘true positive’). In the Results section we report the corresponding averages over N = 20 subjects.

Given the substantial anatomical differences of our subjects, the between-subjects approach required additional pre-processing steps: The coil positions and orientations had to be ‘normalised’. As shown in detail in (Jin et al., 2023), *i*=1, …, *N* positional data 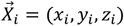 of individual subjects can be readily compared by mapping them on a template MRI yielding 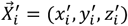. The mapping 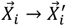 can be defined through warping the cortex-surface of subject-specific MRI to that of the template; we used the MNI152 template (Tadel et al., 2011) and the FreeSurfer-based pial surface (Fischl, 2012). A similar mapping of the coil orientation given as rotation matrix *R* can also be found, at least to some approximation. For this, we estimated the best-fitting affine transform *A* between 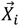 and 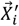 for all *i*=1, …, *N* and used its rotational component *A*rot to transform *R* ′ =*A*rot*R*. We illustrate this for a single subject in Figure 3. The positional data 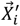 were expressed in spherical coordinates and the rotation matrix *R* ′in Euler angles prior to entering the training/validation/testing procedures.

**Figure 3.**
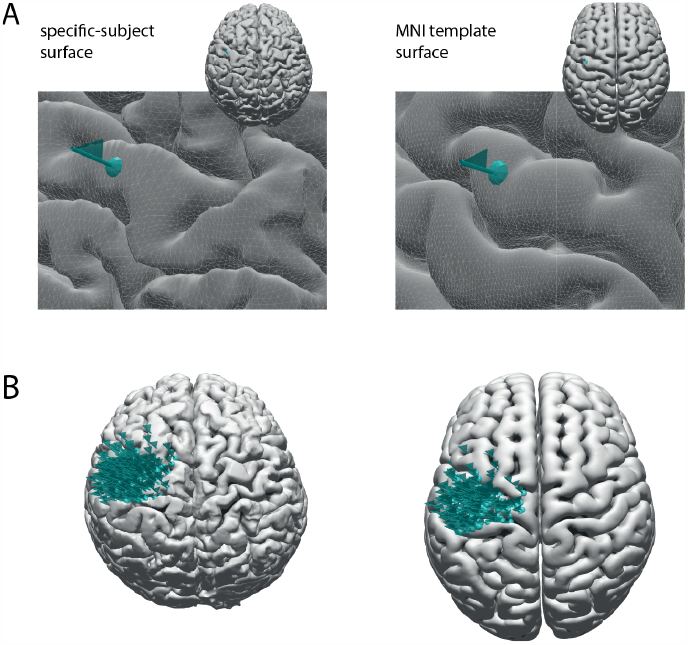
Illustration of the warping position and orientation of stimulations from a single subject’s surface (A) to the MNI template surface (B). The top row details the mapping for a single stimulation whilst the bottom row shows the pattern of all stimulations (here N = 237). The orientation of the flag indicates the TMS coil handle’s opposite direction, while the turquoise dot shows the position of the stimulation.

## 3. Results

### 3.1. Input and response data

We collected 13,851 stimulations from 20 subjects, with an average count of 693 per subject, ranging from 581 to 732; some stimulations were excluded from the current analysis because they accidentally targeted subcortical regions or the ipsilateral hemisphere. Of the included stimulations about 57% (= 7,896) successfully elicited MEPs in either or several of the eight recorded muscles. For the individual subjects this varied between 37% and 84% (average = 395, range = 258 to 570) as shown in Figure 4.

**Figure 4.**
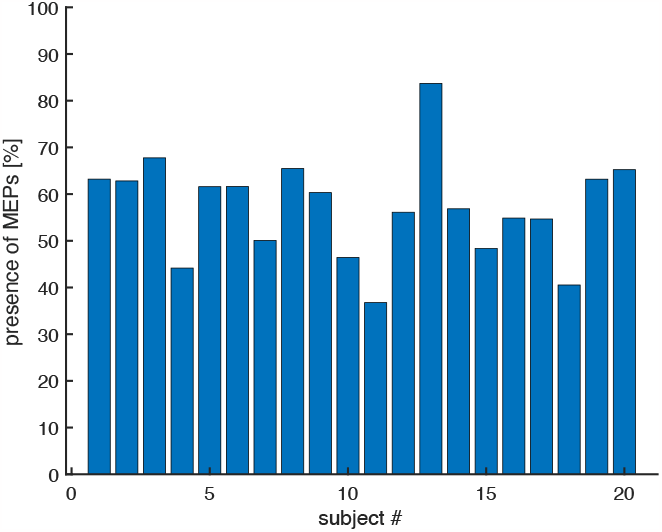
Numbers of successfully elicited MEPs relative to the total number of stimulations per subject. The percentages vary from 37% (subject 11) to 84% (subject 13) readily indicating a sub-stantial between-subject variability in responding to TMS.

### 3.2. Within-subject design

Figure 5 depicts the averaged confusion matrices of the four different machine learners trained and tested in the within-subject design. For all, the predictive capacity exceeded 70%, with the bagging ensemble models showing the maximum performance (77% accuracy). The arguably simple logistic regression model almost reached the same accuracy level (76%), while the more involved support vector machines performed worse (74% for the linear version and 71% for the nonlinear extension via Gaussian kernels). The comparably poor performance of the kernel SVM came as a surprise given that it entails more degrees-of-freedom than its linear counterpart, let alone more than the logistic regression model.

**Figure 5.**
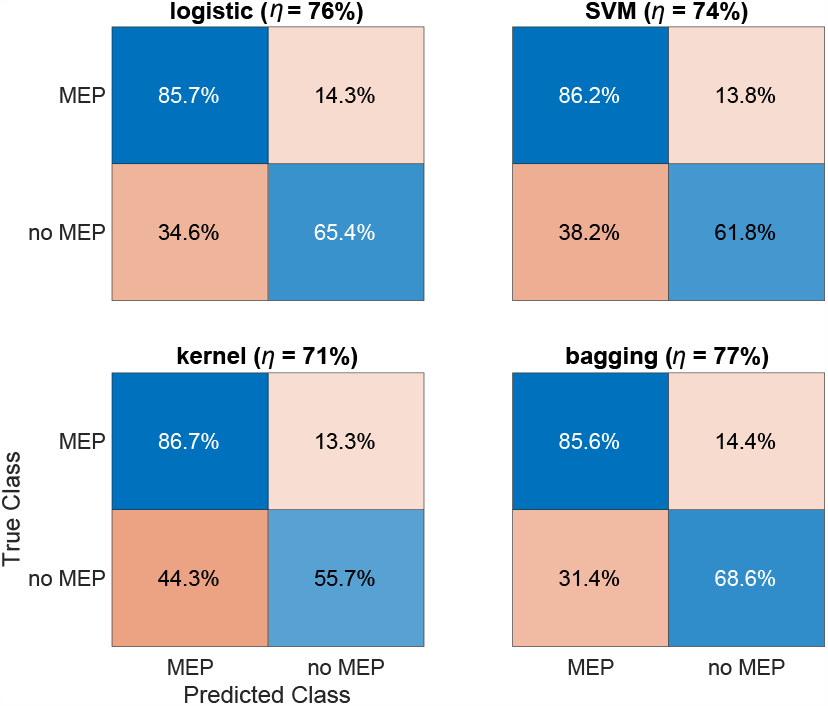
The confusion matrices obtained for the within-subject design. The upper left rectangles (blue) represent ‘true positives’, i.e. prediction of MEPs if they were present, the pink lower left ones indicate false positives (predicting MEPs when non was elicited). The upper right pink rectangles indicate false negatives (predicting no MEPs while MEPs were elicited) and the lower right blue ones represent true negatives. The bagging ensemble approach (lower right panel) displayed maximum performance; see text for details about estimating the model accuracy.

We also generated the confusion matrices per subject from the within-subject design and show this in the Appendix in Figure B1. As expected, we found a substantial between-subject variability that largely matched the variability observed in Figure 4. The maximum accuracy reached 90% while the minimum was a low as 54%, i.e. almost prediction by chance. Since these discrepancies surprised us, we estimated the predictor importance in the case of the tree-based bagging ensemble for all individual subjects and show the corresponding averages in Figure 6; results of the individuals can be found in the Appendix in Figure E1. The predictor importance differed significantly across subjects which possibly influenced results when using the between-subject design.

**Figure 6.**
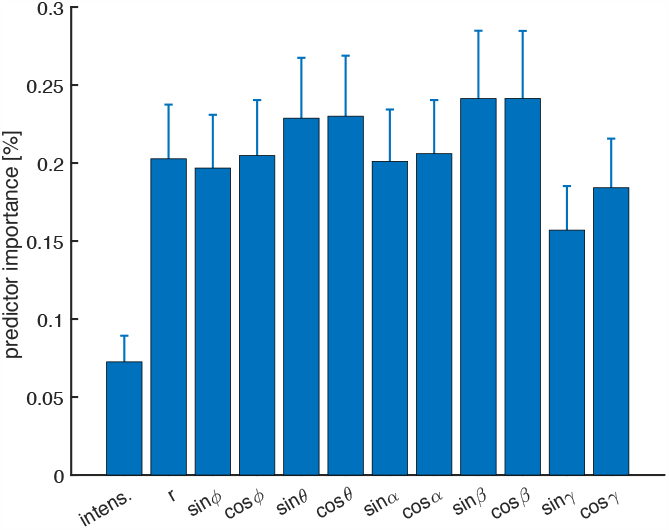
Mean importance for each of the twelve predictors. We computed accumulating changes in the node risk due to splits in the tree, and then dividing the sum by the total number of branch nodes. There the change in node risk is the difference between the risk for the parent node and the total risk for the two children. Error bars indicate the standard error over subjects.

### 3.3. Between-subject model

The performance in the between-subject design was clearly less than in the within-subject design. Again, the bagging ensemble turned out to be best, but did only reach a predictive capacity of about 65%, which was very similar to the logistic regression, as shown in Figure 7. Again, both SVM approaches performed less although the conventional SVM displayed the highest true positives prediction (almost 88%; see top right panel in Figure 7). The corresponding confusion matrices for each subject in the between-subject design can be found as Figure B2 in the Appendix. The maximum prediction accuracy reached a maximum of 72%, i.e., it was clearly lower than when training any of the models in the within-subject design.

**Figure 7.**
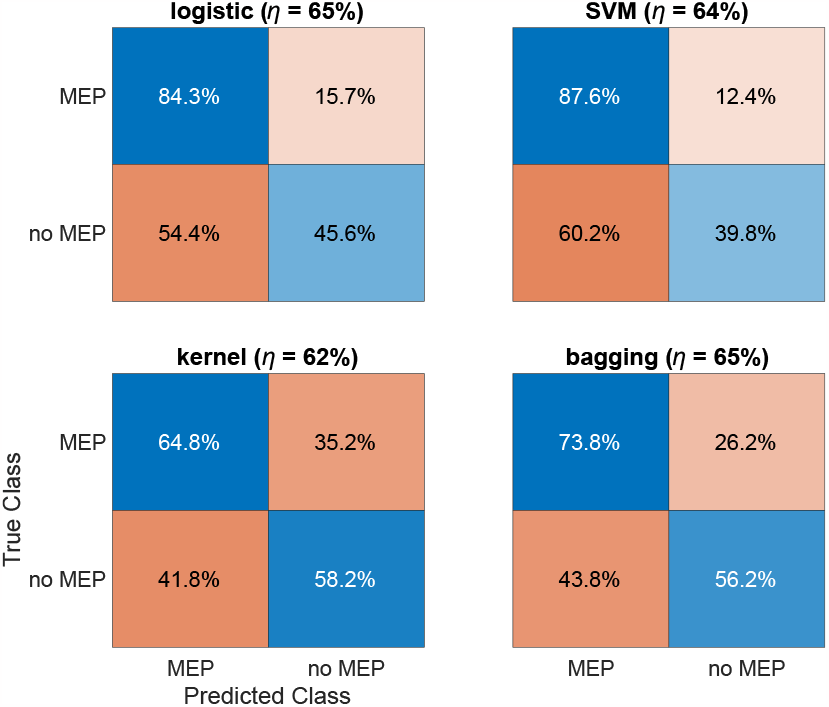
The confusion matrices obtained from the betweensubject design; cf. Figure 5. Again, the bagging ensemble approach yielded on average the maximum accuracy, but performance clearly deteriorated when compared to the within-subject design arguably due to the pronounced between-subject differences despite standardising data to the MNI template MRI.

## 4. Discussion

We predicted the presence of MEPs using different machine learning approaches. The machine learners were either trained separately on every of the N = 20 participating subjects (within-subject design) or via hold-one-out partitioning, i.e. training using data of N-1 = 19 subjects and testing on the remainder (between-subject design). While the within-subject design yielded acceptable prediction accuracy (acceptable since the amount of data for training was limited), the results of the between-subject design were disappointing (here the amount of data for training was about 20 times larger and deemed sufficient).

For both within and between subject designs, the decision tree approach combined as a bagging ensemble showed the maximum performance. This is not surprising as this machine learner is known for its rich predictive capacity and – when it comes to boosting alternatives (see around Table A1), it is considered particularly robust against random variability that we observed in our data (see Figures 4 & 6 as well as C1 & E1 in the Appendix). Interestingly though, the performance gain over, e.g., the arguably simple logistic ridge regression, was not spectacular, suggesting that either the sample size did not suffice to train the ensemble well enough or that the data really did not contain sufficient information to make the desired predictions.

The performances of (most of) the subject-specific models speak against the latter. This brings us to potential limitations when combining our data to suit for between-subject comparisons. Essential here is the warping of the subject-specific anatomy to a template head. While this is common practice in, e.g., MRI-assessments, integrating the concomitant transforms in TMS is rarely done; see (Jin et al., 2023) for an exception. We chose to define the transform by matching pial rather than grey matter surfaces. This has the advantage to not suggest spatial resolutions that TMS cannot provide, it does limits estimates of ‘really’ excitable areas in the cortex. In a way this limitation suggests replacing this pre-processing step by proper estimates of the induced electric fields, though when doing so, one must extract selected scalar values of these fields to-be-used as predictors in a machine learning approach. Unlike previous work, here, we not only transformed the focal point of the magnetic field, i.e., its position, but also its orientation. We follow a ‘poor man’s’ approach and simply estimated an affine transform based on the original and transformed positional data. While this guaranteed the smoothness of the transform, it ignored that warping may, for example, cause a point that was originally on top of a gyrus to end-up close to a sulcus. In that case the change in curvature may be substantial and may require more than the global affine transform to ‘correct’ coil orientation. Implementing such a more local transform comes with challenges as only two of the three orientational angles are uniquely defined. Here we tested for the possibly deteriorating effect of transforming the data to MNI-space by comparing the results in Figure 5 that were determined using the original subject-specific MRIs with those computed on the MNI-template following the within-subject design. Figures D1/2 clearly show that transforming TMS stimulation points to a template hardly comes with information loss and, consequently, does not lead to a reduced prediction accuracy of our machine learners. Given the challenges in transforming coil orientation one may wonder whether one may ignore it altogether when seeking for prediction models for TMS. We tested this by rerunning our machine learning procedures without any information about coil orientation. As shown in Figures F1/2, it is appropriate to the include coil orientation to attain (slightly) better prediction accuracy. This is consistent with earlier findings about the relevance of TMS orientation when eliciting MEPs (Opitz et al., 2013; Vink et al. (2018).

As said, we are disappointed by the low accuracy of the between-subject models. The substantial individual differences in MEPs occurrence may be seen as a possible cause for the limited performance (cf. Figure 4). The pronounced differences in predictor importance between subjects supports this impression (cf. Figure 6). By the same token current (non-linear) machine learners should have the capacity to cope with this variability provided the input data do contain sufficient information to form the to-be-identified classes and to train the classifier. While incorporating more than 10,000 data points when training our model sounds like a lot, we must admit that they stemmed from only 19 subjects. In previous studies, predictions using machine learning were made using a larger set of training data. For example, Zheng et al. (2015) aimed to identify Alzheimer’s disease (AD) and mild cognitive impairment (MCI) in subjects by using the connectivity networks of MRI to differentiate and diagnose these diseases. They trained a machine learning model on MRI data from 163 patients with Alzheimer’s disease and 189 patients with MCI, and they achieved accuracies of 92.1% and 86.9%, respectively. Previous research with smaller subject numbers produced much lower accuracies. Dayan et al. (2018) employed machine learning to classify large- or small-amplitude MEP classes using the cortical structural characteristics of MRI. Based on data from 25 subject, the accuracy of the MEP categorisation did not exceed 66.67%. Although there the research question and input data differed entirely from ours, it suggests that with a sufficiently large sample size, the between-subject variability might become of minor importance when predicting scalar, categorical outcome measures.

While the results for the within-subject design are more encouraging, also in this case there is room for improvement. For example, one may add additional features to the original input data, which in our study only consisted of the position, orientation, and intensity of TMS. For instance, Stokes et al. (2007) reported that the motor threshold rose with the coil-cortex distance, i.e., that distance could be one of these additional features. Moreover, the individual hand-knob curvature also correlates with the presence of MEPs Opitz et al. (2013). Future studies may consider further features including morphological characteristics of the subject’s MRI (as, e.g., in Dayan et al. 2018) as such one can expect most such additions have the potential to improve the predictability of MEPs when added as input in a machine learner.

## 5. Conclusion

We used the TMS intensity and coil position and orientation to predict the presence of MEPs in a group of hand and forearm muscles. Compared to more traditional machine learners, we achieve maximum higher accuracy when using decision tree-based bagging ensembles, especially when adding coil orientation to the set of predictors. Predicting TMS response within a subject appears in reach of machine learning approaches despite the limited sample size. Predicting a subject’s responses by training machine learners with data of other subjects, however, remains a challenge.

## Acknowledgements

FJ would like to thank the Chinese Scholarship Council for financial support [201706210060]. SMB was funded by a VIDI grant from the Dutch Research Council [016.Vidi.178.014].

## Appendix

*Models and hyperparameters*

We determined the hyperparameters a priori pooling all data. This initial optimisation step selected for the logistic regression model and for the linear SVM a ridge regression over a L1 regression (LASSO) for regularisation and for the non-linear SVM Gaussian kernels over radial basis functions. For the bagging example the random decision tree approach with Gini’s diversity index as split criterion outperformed adaptive (logistic), gentle, robust, random undersampling, and totally corrective boosting; see Table A1 for the result.

**Table A1.** Matlab functions and hyperparameters for the four machine learning models in the between-subject design.

| model | function | hyperparameter | value | hyperparameter | value |
| --- | --- | --- | --- | --- | --- |
| Logistic regression | fitclinear | Score transform | logit | Beta | -41.56 – 8.84 |
| | | Lambda | $1.46 \cdot 10^{-7}$ | Beta tolerance | $10^{-4}$ |
| | | Bias | 68.77 | Gradient tolerance | $10^{-6}$ |
|  |  | Regularisation | ridge | Iteration limit | 1,000 |
| SVM | fitclinear | Score transform | logit | Beta | -25.00 – 8.83 |
| | | Lambda | $2.19 \cdot 10^{-8}$ | Beta tolerance | $10^{-4}$ |
| | | Bias | 64.95 | Gradient tolerance | $10^{-6}$ |
|  |  | Regularisation | ridge | Iteration limit | 1,000 |
| kernel SVM | fitcsvm | Score transform | logit | Kernel scale | 0.14 |
|  |  | Kernel function | Gaussian | Solver | SMO |
| bagging ensemble | fitcensemble | Score Transform | logit | # learning cycles | 482 |
|  |  | Learner | binary classification decision tree with surrogate splits |  |  |
|  |  | Split criterion | Gini's diversity index (gdi) |  |  |
|  |  | Min leaf size | 1 | Max # splits | 11051 |
|  |  | Score Transform | logit | Min leaf size | 1 |

The hyperparameters varied slightly for consecutive estimates due to randomisation of initial conditions though the resulting k-fold loss hardly differed, and for the subject-specific model they largely agreed with the ones of the pooled data shown in Table S1.

**Figure B1.**
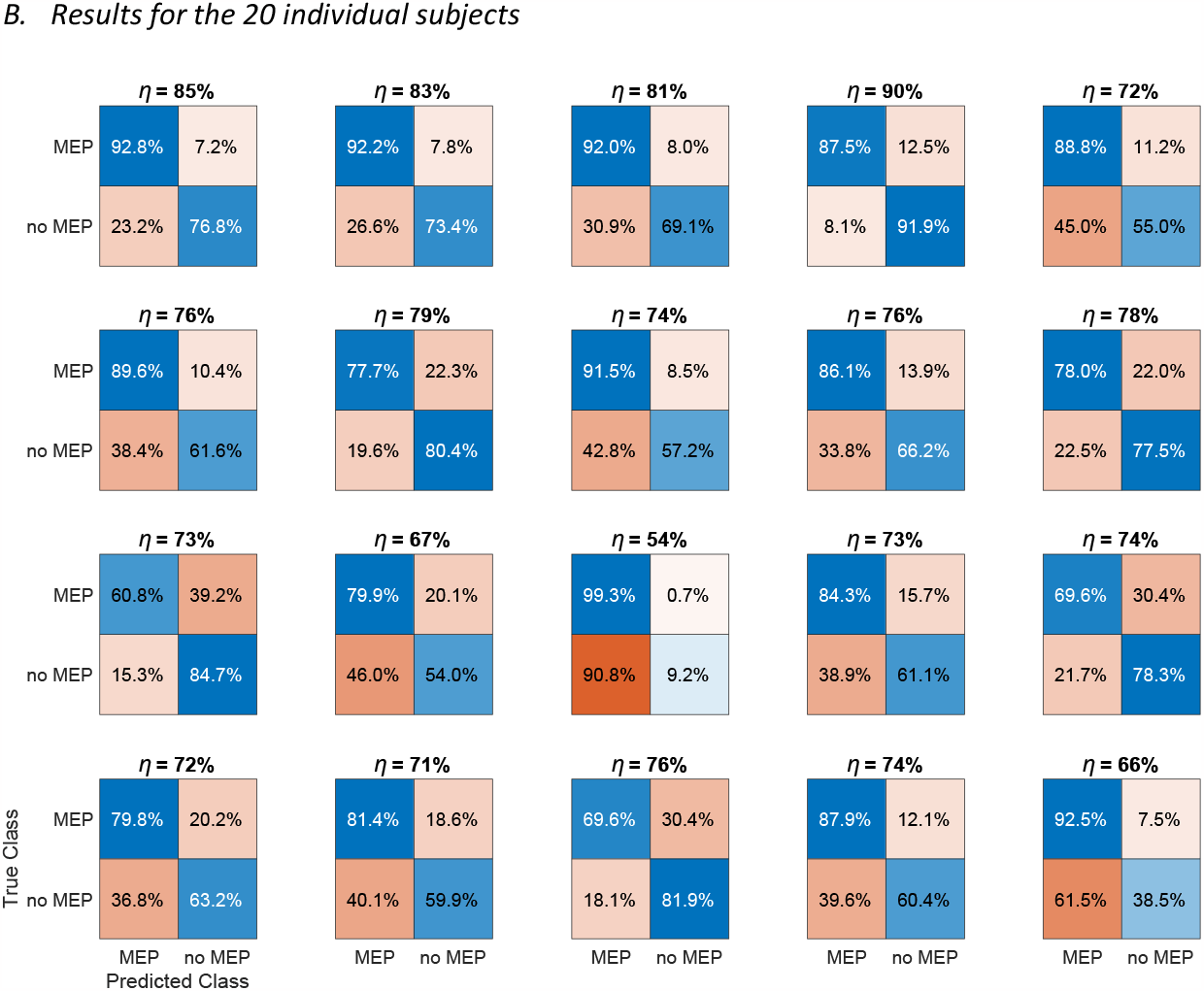
The results for the within-subject design shown in Figure 5 in the main text but split up for the individual subjects. Please note the presences of outliers like subject 13, where the accuracy is as low as about 56%; cf. Figure 4 in the main text.

**Figure B2.**
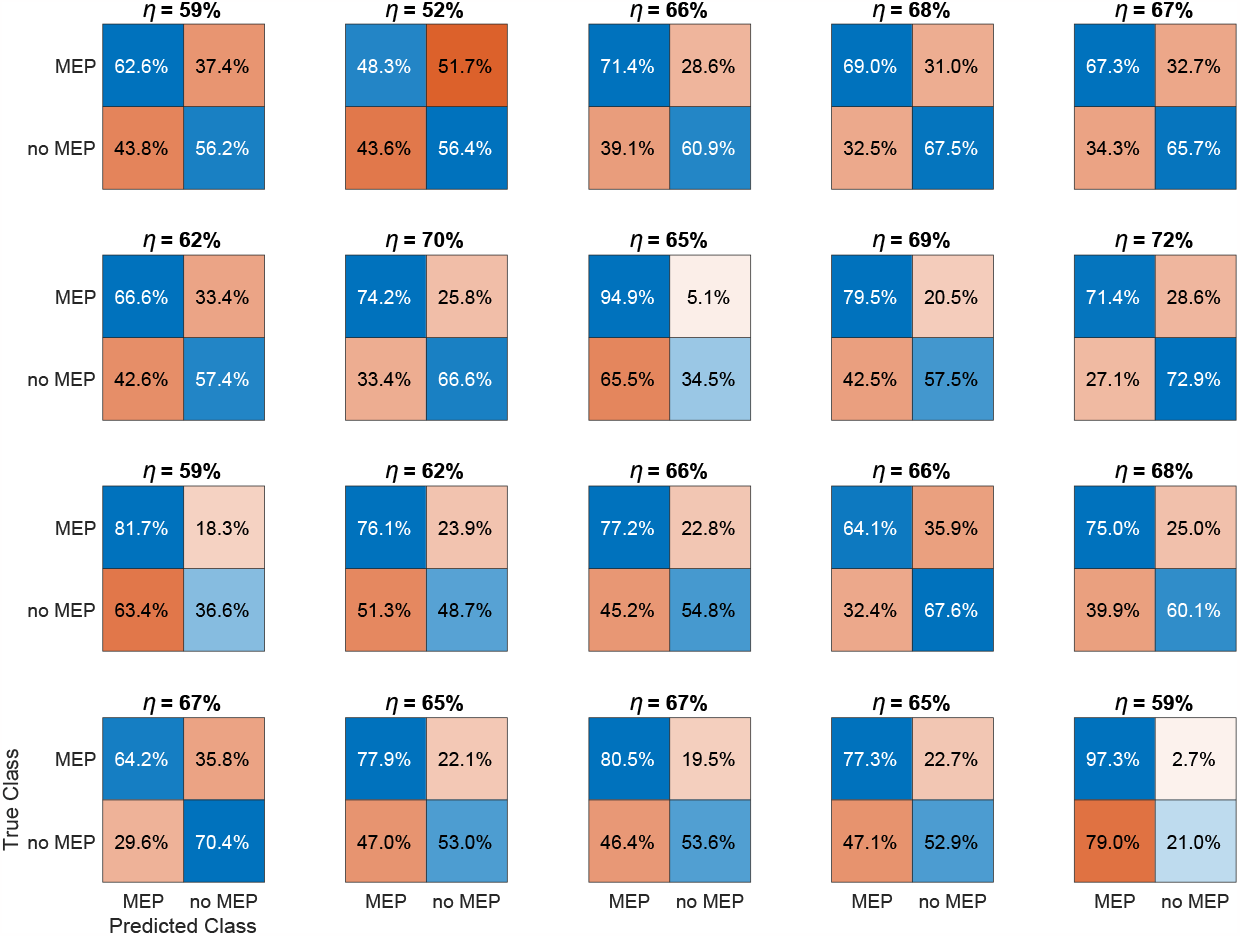
The results for the between-subject design shown in Figure 7 in the main text but split up for the individual subjects.

**Figure C1.**
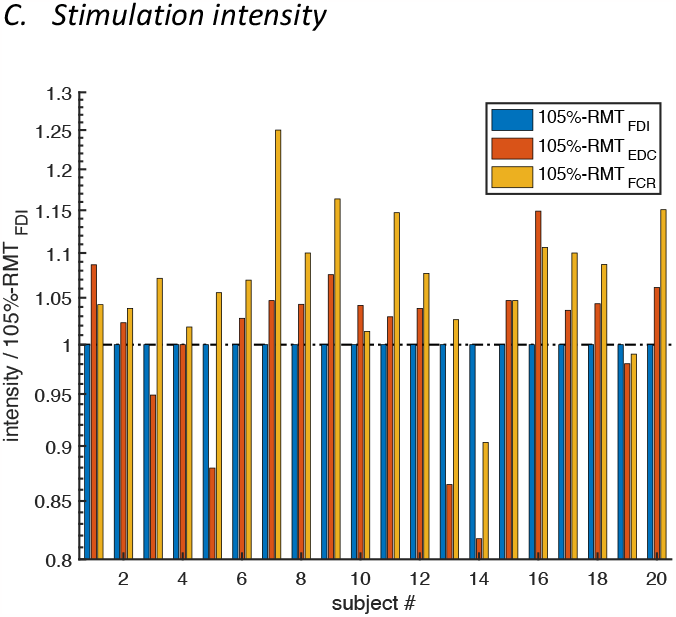
Stimulation intensities per subject relative to the subject-specific 105%RMT_FDI_. Like the number elicited MEPs shown in Figure 4, also the great differences in RMT indicate a substantial between-subject variability in responding to TMS.

**Figure D1.**
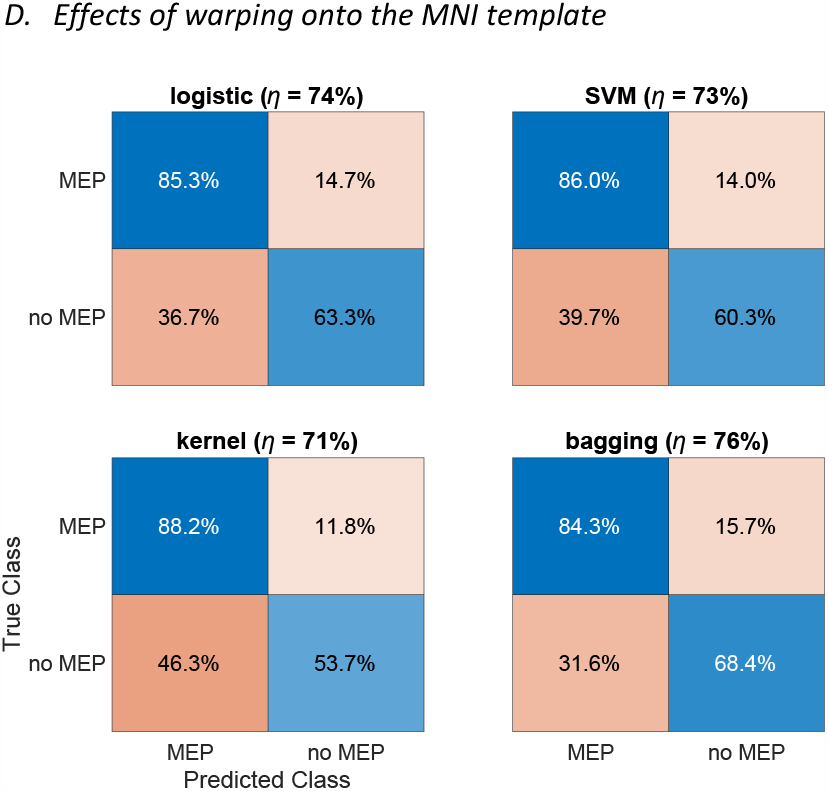
Same as Figure 5 but warping the data onto the MNI template.

**Figure D2.**
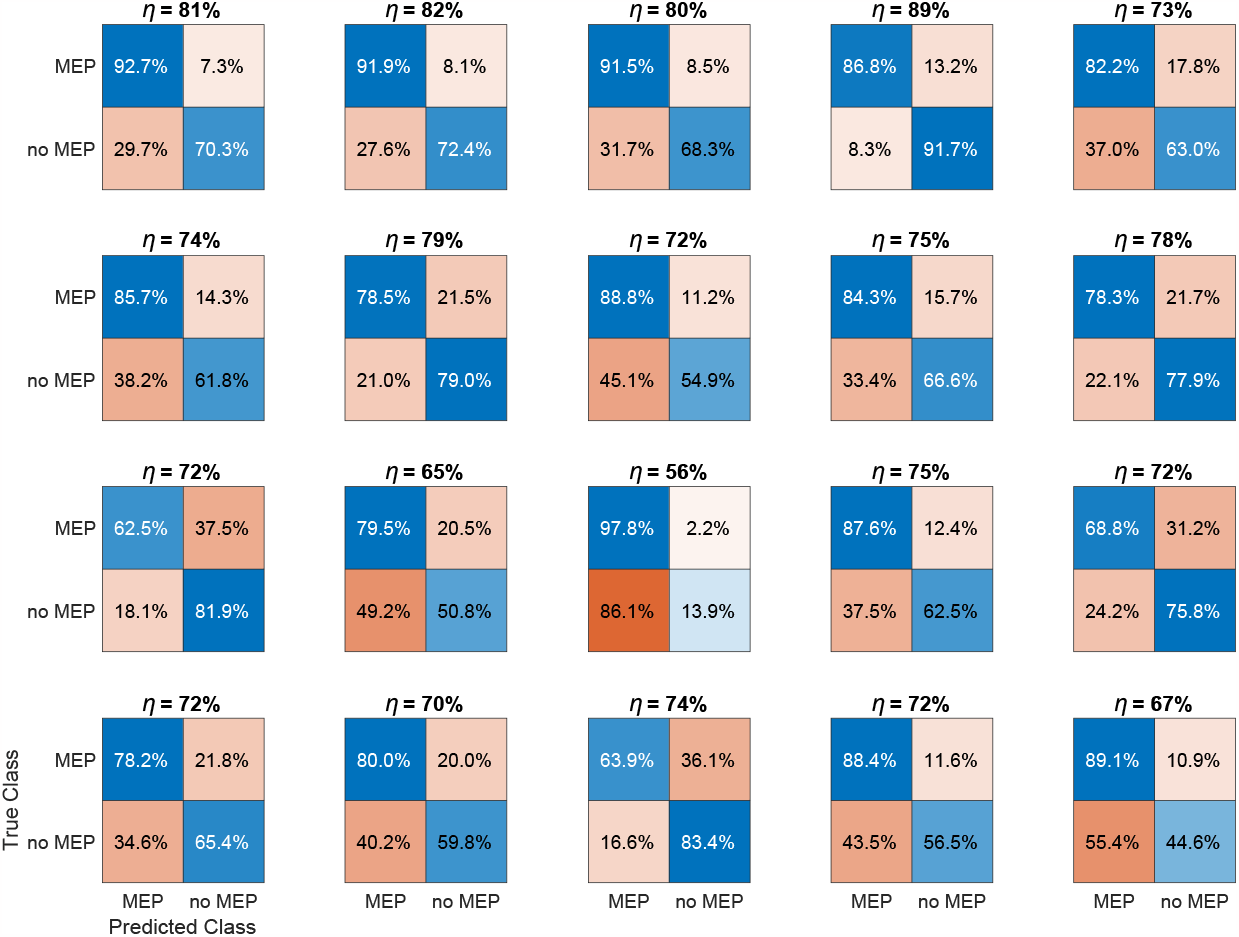
Same as Figure D1 but split up for the individual subjects; compare also Figure B1.

**Figure E1.**
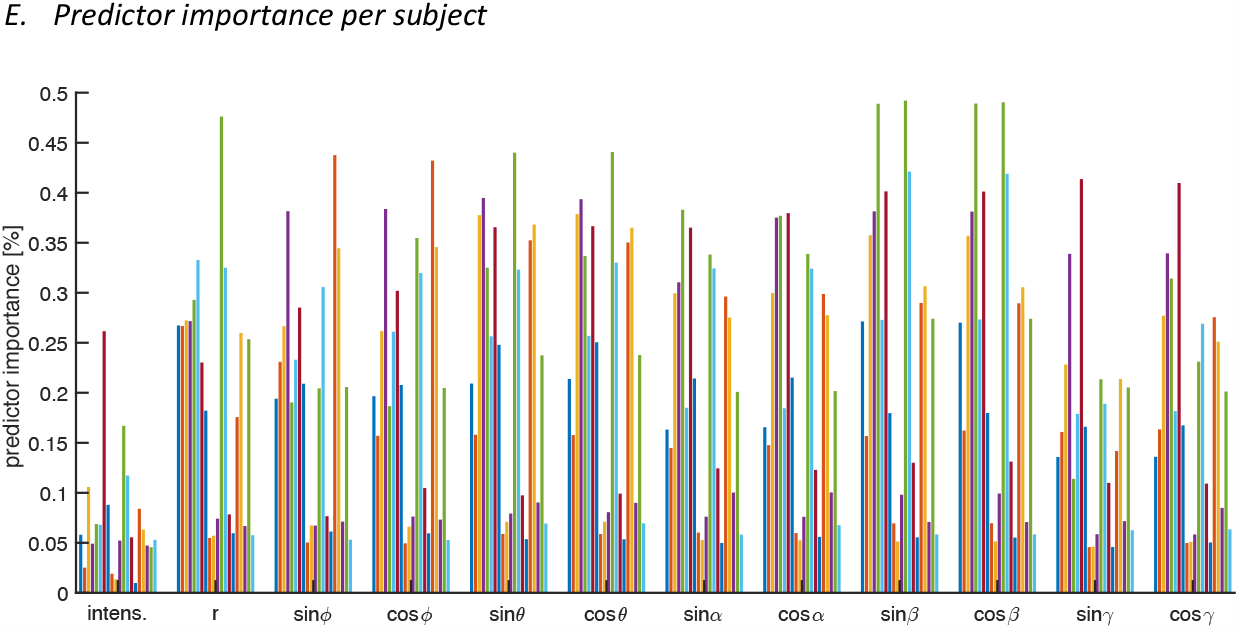
Predictor importance for the 20 individual subjects; see main text for further explanation.

**Figure F1.**
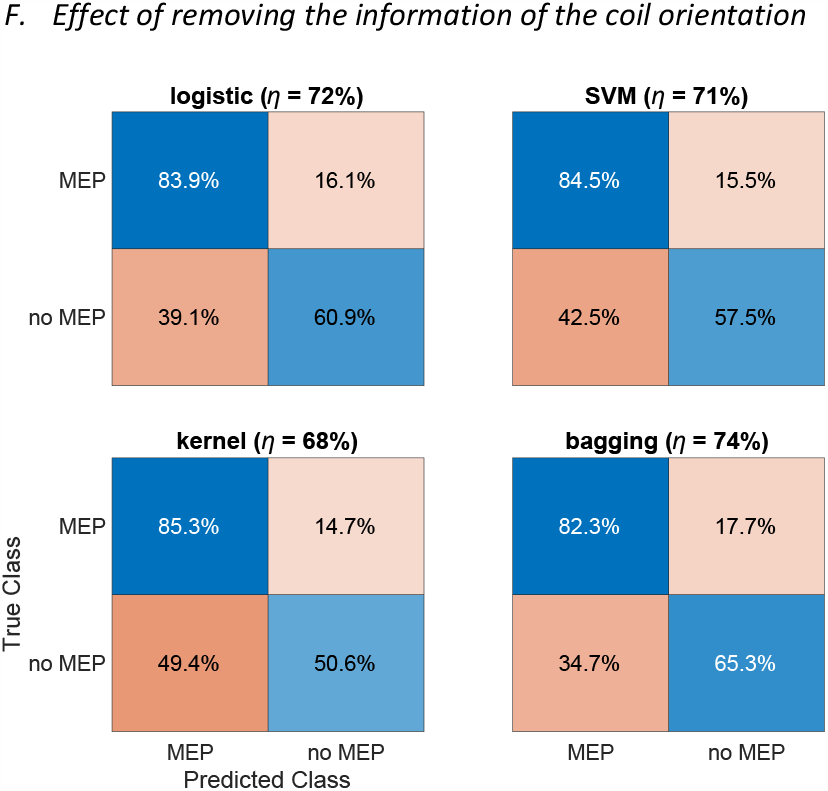
Same as Figure 5 but without including the information about the coil orientation (Euler angles). Performance drops slightly.

**Figure F2.**
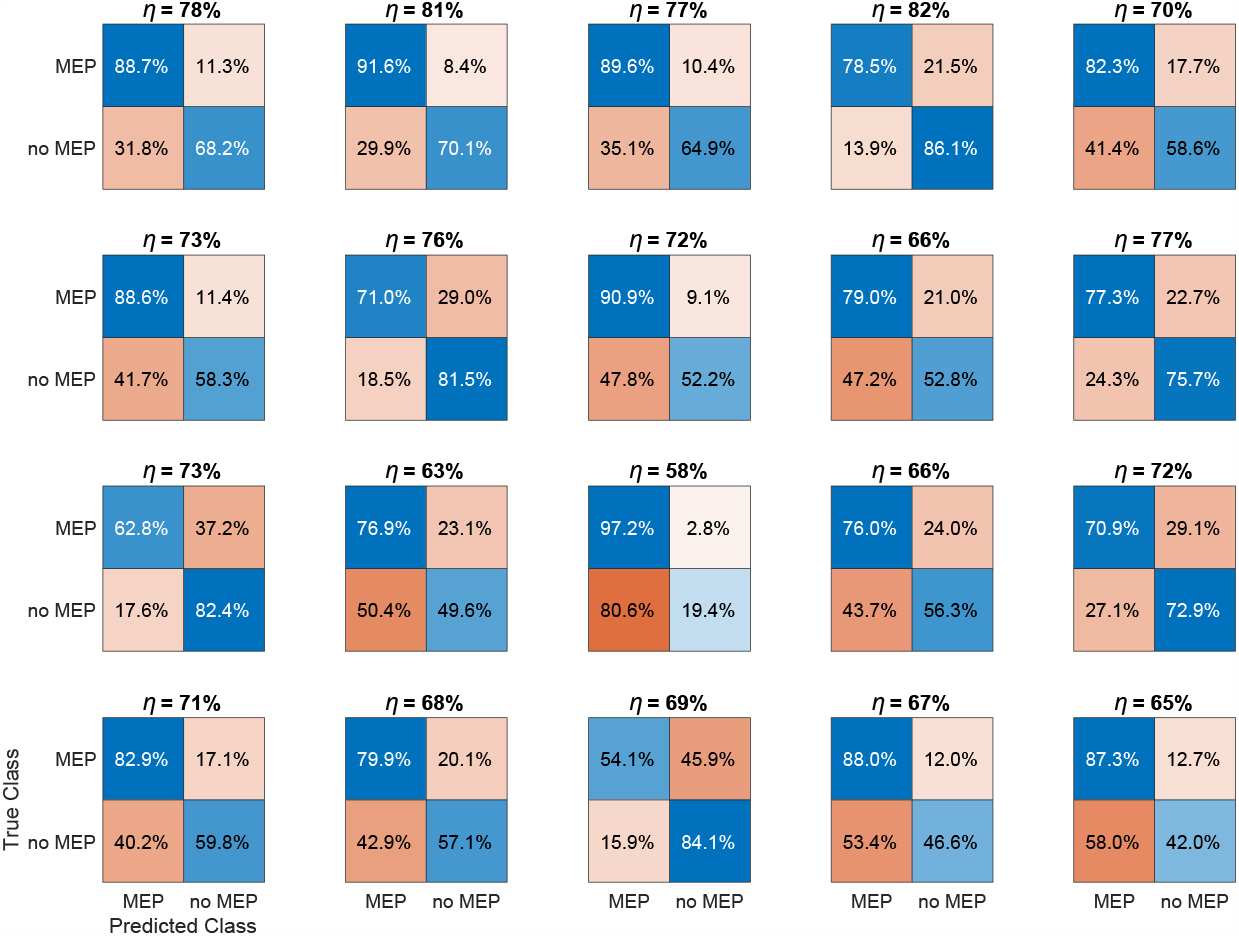
Same as Figure F1 but split up for the individual subjects; compare also Figure B1.

